# Synthetic Rewiring of Virus-Like Particles via Circular Permutation Enables Modular Peptide Display and Protein Encapsulation

**DOI:** 10.1101/2025.06.13.659512

**Authors:** Shiqi Liang, Kaavya Butaney, Daniel de Castro Assumpção, James Jung, Nolan W. Kennedy, Danielle Tullman-Ercek

## Abstract

Virus-like particles (VLPs) are self-assembling nanoparticles derived from viruses with potential as scaffolds for myriad applications. They are also excellent testbeds for engineering protein superstructures. Engineers often employ techniques such as amino acid substitutions and insertions/deletions. Yet evolution also utilizes circular permutation, a powerful natural strategy that has not been fully explored in engineering self-assembling protein nanoparticles. Here, we demonstrate this technique using the MS2 VLP as a model self-assembling, proteinaceous nanoparticle. We constructed, for the first time, a comprehensive circular permutation library of the fused MS2 coat protein dimer construct. The strategy revealed new terminal locations, validated via cryo-electron microscopy, that enabled C-terminal peptide tagging and led to a stable protein encapsulation strategy via covalent bonding – a feature the native coat protein does not permit. Our systematic study demonstrates the power of circular permutation for engineering new features as well as quantitatively and systematically exploring VLP structural determinants.

## Introduction

Virus-like particles (VLPs) are self-assembling protein nanoparticles adapted from viruses by removing their replication and infection components. VLPs typically assemble spontaneously from multiple copies of one or more viral coat proteins (CPs) and exhibit high stability and biocompatibility. Their uniform size and defined chemistry make them promising scaffolds for several applications, including imaging, vaccination, drug delivery, and as biomaterials^1–3^. To fully reach their potential in these areas, several key features of VLPs need to be engineered, including encapsulation of target cargo as well as decoration of the particle surface to specific targeting ligands or antigens. Protein engineering on viral CPs provides a way to stably introduce peptide tags and conjugation-compatible amino acids into VLPs, permitting incorporation of those new features into the naturally evolved proteinaceous nanoparticles^4,5^. To date, synthetic biologists have adopted several genetic and protein engineering approaches borrowed from natural evolution, including amino acid substitutions, short sequence insertions, and gene duplication/fusion^4–10^. However, in nature, evolution utilizes another complicated strategy to diversify protein sequence and structure prior to the selection, which has not been fully explored in engineering self-assembling protein nanoparticles – circular permutation^8^.

Circular permutation is a naturally occurring strategy to enlarge the protein sequence and structure space^11^. This process connects the original N- and C-termini of a protein and relocates the termini to a new location while retaining the overall protein structure^12^. In naturally evolved proteinaceous assemblies, there are cases where protein subunits are circularly permuted to adopt novel functions^13,14^. This strategy has also been applied to engineer natural self-assembling protein systems for diverse functions, such as modifications at new termini or novel assembly morphologies^15–19^. However, those circularly permuted proteins were largely based on rational design and to date there is no systematic circular permutation study on a self-assembling proteinaceous nanoparticle to fully explore the sequence and structure space of the subunit protein. In this study, we sought to systematically assess the tolerance to circular permutation of a model self-assembling protein system, the *Escherichia coli*-infecting Male-Specific 2 (MS2) VLP, with the goal of engineering novel capabilities.

Derived from the MS2 bacteriophage, the wildtype (WT) MS2 VLP is composed of 180 MS2 CP copies and adopts an icosahedral T = 3 structure with a diameter of 26-28 nm^20^. During the assembly process, MS2 CPs first dimerize, and these CP dimers function as basic building blocks forming the full VLP.^21^ CP dimers adopt two different conformations in assembled MS2 VLPs – 30 in the symmetric C/C conformation and 60 in the asymmetric A/B conformation (Fig. 1)^20,22^. The main difference among the three different CP conformations is at the FG-loop that mediates subunit interactions within VLP facets^20,22^. A- and C-conformers have extended FG-loops pointing to pores at the hexagonal facets^20,22^. However, the FG-loop of B-conformer adopts a compact conformation, mediating the interaction at the pores of the pentagonal facets^20,22^. The interior surface of MS2 protein shell is positively charged, enabling it to encapsulate RNA cargo during the assembly process^22–25^. The encapsulated cargo can be expanded beyond RNA to other molecules with negative charges, including DNA^26^, nucleic acid conjugates^27,28^, and proteins with anionic tags^26^.

**Figure 1.**
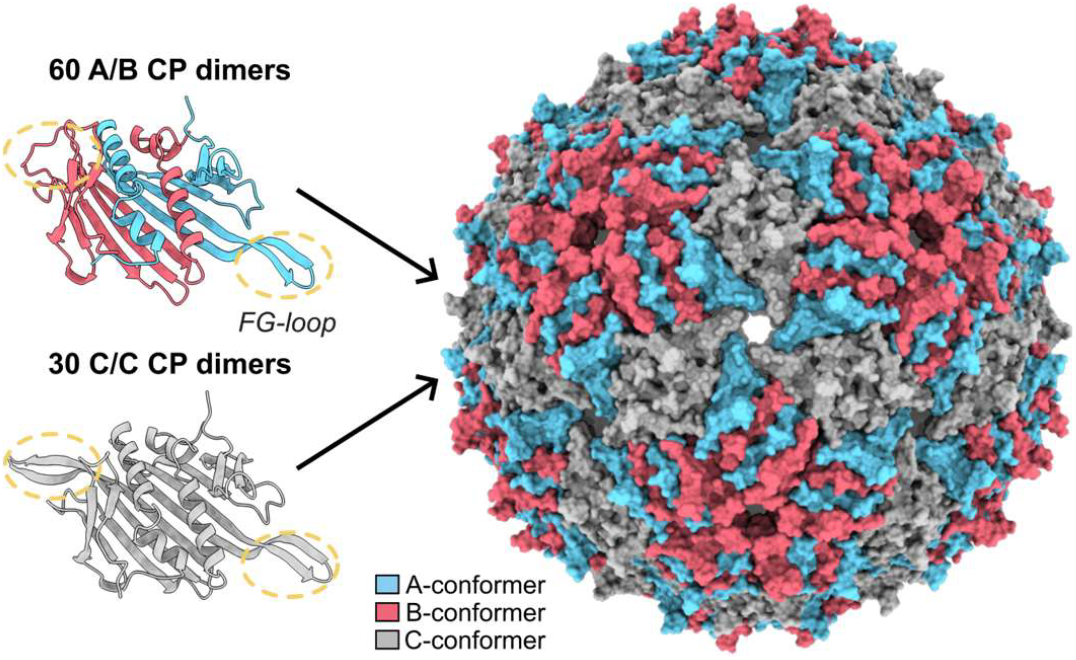
Protein composition of the MS2 VLP. Spontaneously formed MS2 CP dimers adopt 2 different conformations in the assembled VLP. A/B-conformers build up pentagonal facets with compact FG-loops (circled with yellow dashes) of B-conformers pointing towards pores. In the hexagonal facets, FG-loops of A- and C-conformers are in extended conformations and mediate interactions between subunits. (PDB ID: 1ZDI)

The MS2 VLP is a great model to systematically study self-assembling proteinaceous nanoparticles. One advantage provided by this system is its ability to encapsulate RNA molecules when expressed in host cells, building a phenotype-genotype linkage in assembled VLPs. Based on this property, the Systematic Mutation and Assembled Particle Selection (SyMAPS) method enables systemic engineering of VLPs and study the effect of modifications on VLP assembly^9^. The method was applied to the MS2 CP to study the effect of single mutations^9^, epistatic residue mutations^29^, and random short peptide insertions to the N-terminus^5^ and in the FG-loop^10^. Those studies identified interactions between specific residues that are essential for VLP assembly. However, mutational scans and loop insertions reveal little about the essentiality of CP structure elements because these types of modifications rarely have large effects on the CP backbone. Understanding the influence of larger changes on CP folding and VLP assembly is needed to facilitate adding foreign peptides on the exterior or interior surface of VLPs to enable enhanced delivery^30^, antigen display^31,32^, and enzyme encapsulation^33^. However, genetically attaching foreign peptides to the MS2 CP without disrupting VLP assembly still remains challenging, as adding tags to either the N- or C-termini and inserting peptides into the CP typically leads to heterogeneity in VLP morphology or failure to assemble^5,10,31,33,34^.

Here, we created a library of all possible circular permutations on a genetically fused MS2 CP dimer to alter the terminal locations and create potentially accessible loci for peptide fusion. We generated a comprehensive circular permutation apparent fitness landscape (AFL) of MS2 CP using SyMAPS and characterized the assembly competency of all circular permutants. Our circular permutation AFL quantitatively describes the effect of disrupting original structure elements in the MS2 CP folding and VLP self-assembly process. Moreover, our systematic screening identifies positions tolerant to introducing new termini, which enables terminal peptide tagging and, in turn, a new protein encapsulation strategy via covalent bonding. Our study explores the MS2 CP sequence and structure space for VLP self-assembly and provides new MS2 VLP engineering platforms with assembly-competent permutants. Additionally, we demonstrated the utility and feasibility of adopting a systematic approach toward leveraging circular permutation for engineering novel functions, which could be expanded to many other VLP systems.

## Results

### Generating a circular permutation library based on MS2 CP-CP

We sought to use circular permutation to create new N- and C-termini of the MS2 CP to circumvent challenges with peptide display on the WT protein. Circular permutation relies on covalently connecting the N- and C-termini of the protein^12^. Therefore, the distance between protein N- and C-termini is often a key factor for the performance of a circularly permuted protein target. With this in mind, we chose to use a fused CP dimer (CP-CP) as our scaffold. In this dimer, the C-terminus of the MS2 CP is genetically encoded as a direct fusion to the beginning of a second, identical MS2 CP^6^. Given that this CP-CP fusion has close termini and naturally functions as the protomer in MS2 VLP assembly, we hypothesized that CP-CP would be amenable to circular permutation. We confirmed its ability to assemble into VLPs by size-exclusion chromatography (SEC), sodium dodecyl sulfate-polyacrylamide gel electrophoresis (SDS-PAGE), negative stain transmission electron microscopy (TEM) and dynamic light scattering (DLS) (Supplementary Fig. 1 and Supplementary Table 1). Given that CP-CP does not contain any sequence modification other than duplicating the monomeric CP sequence, the spatially close N- and C-termini of CP-CP make it an excellent scaffold for performing circular permutation. This allows us to test for disruptions to VLP assembly after introducing termini in new locations, disrupting structural elements within the MS2 CP (Fig. 2a).

**Figure 2.**
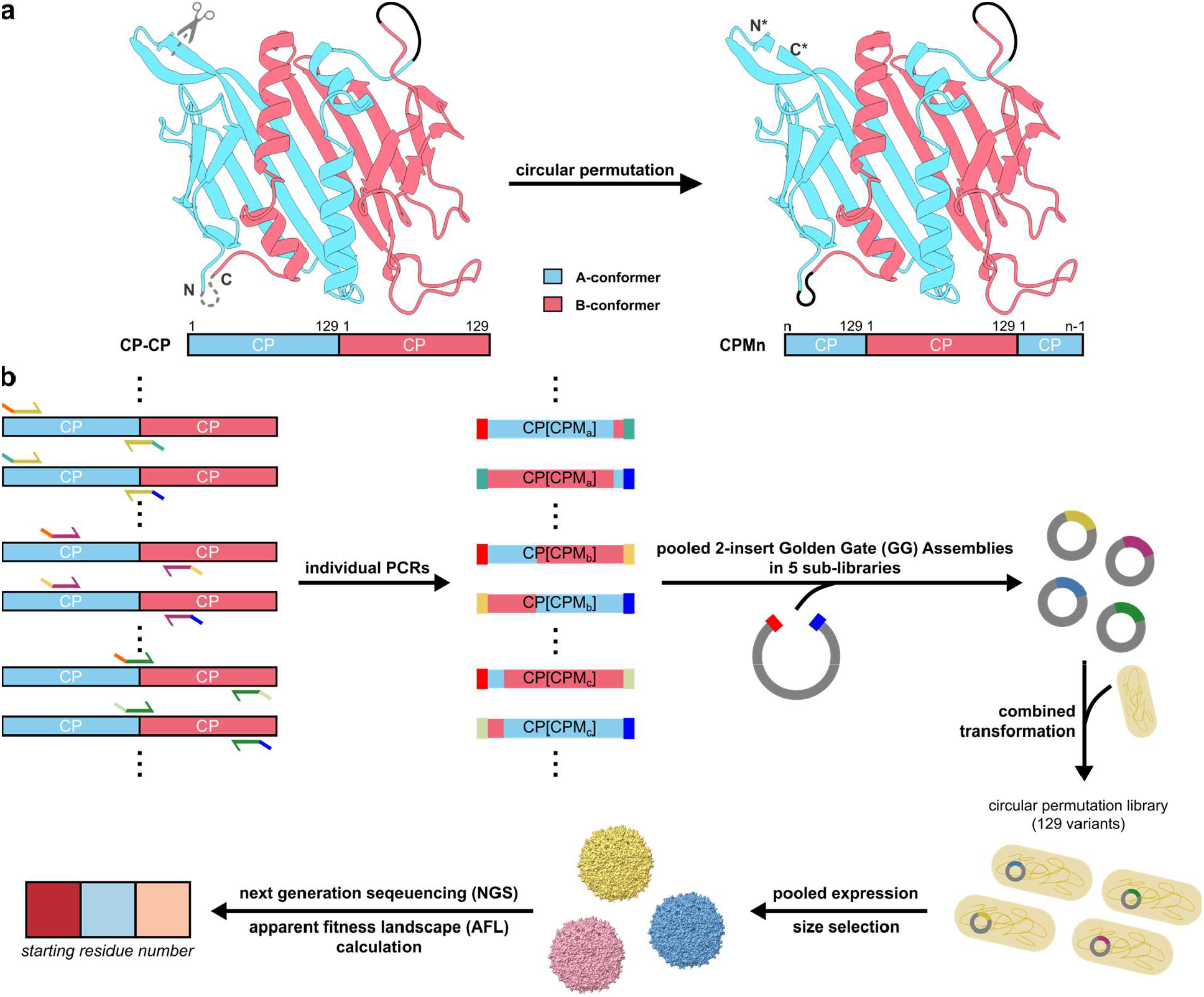
Performing circular permutation on MS2 CP-CP. **a**, Schematic of the approach of performing circular permutation on MS2 CP-CP (PDB ID: 1ZDI). To generate permutant n (CPMn), the first (n-1) residues of CP-CP were moved to the C-terminus. Therefore, CPMn starts at the nth residue of WT CP, followed by an intact CP copy, and ends at the (n-1)th residue of WT CP. **b**, Diagram of generating the circular permutation apparent fitness landscape (AFL). The comprehensive circular permutation library was constructed by pooled two-insert Golden Gate (GG) Assemblies in five sub-libraries, followed by a combined transformation of all 129 variants into Escherichia coli. A size selection enriched VLP-forming permutants and helped to calculate the circular permutation AFL.

To identify locations amenable to the introduction of the new N- and C-termini, we created a comprehensive library covering all 128 possible locations plus the original N- and C-terminus of CP-CP. We utilized a two-insert Golden Gate (GG) assembly strategy to construct the library^35^, where one piece of the GG insert encodes one CP permutant copy. We then expressed the library of variants and purified resulting VLPs via a process that includes a size selection for MS2 VLP-like structures (Fig. 2b). According to the SEC chromatograph, assembled VLPs showed a high absorbance at 260 nm, indicating the encapsulation of genetic information within assembly-competent variants (Supplementary Fig. 2a). The presence of VLPs was confirmed by negative stain TEM (Supplementary Fig. 2b). We expected that only a few permutants would retain assembly competency and survive the size selection step. Indeed, the yield of assembled VLPs was much less than the yield of the unpermuted CP-CP VLPs from an equivalent production scale, suggesting that only a few variants were present after the size selection (Supplementary Fig. 2a).

### AFL reveals circular permutation tolerant sites

Next, we used next generation sequencing (NGS) to quantify the abundance of all variants before and after the size selection. All 128 variants, and the non-permuted CP-CP, were identified in our plasmid library prior to the size selection process, indicating that our library included full coverage of all theoretically possible circular permutants. We compared the relative percent abundance of each variant before and after the size selection across three biological replicates and generated a quantitative AFL with apparent fitness scores (AFSs) and deviations describing assembly competency of all variants (Fig. 3a and Supplementary Data).

**Figure 3.**
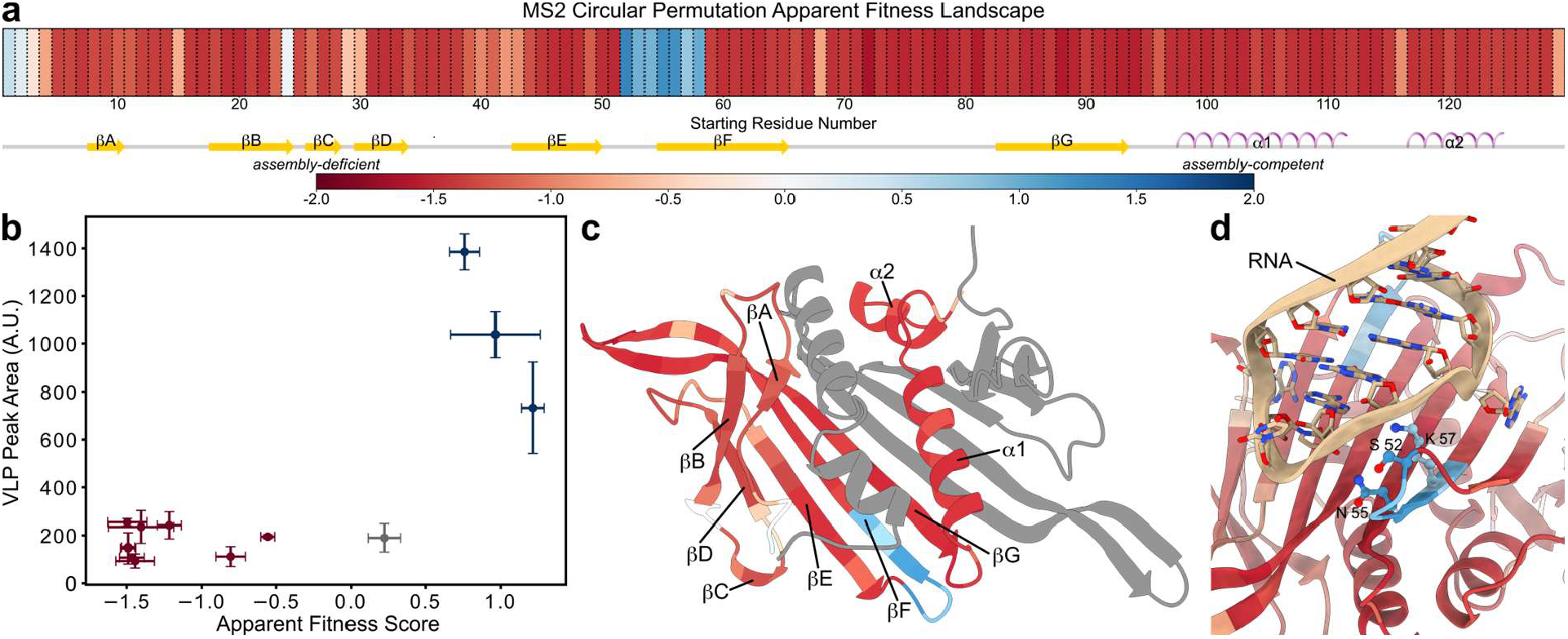
MS2 circular permutation AFL. **a**, Apparent fitness scores (AFS, n = 3) for all permutants starting at diferent residues in the first CP of CP-CP. **b**, Individual assembly assays (n = 3) on 13 randomly selected variants correlate well with AFS values. Blue points were classified as assembly-competent permutants, red points exhibited VLP assembly deficiency, and gray points were identified as intermediate permutants. Error bars indicate standard deviations of assembly quantification and AFS. **c**, C-conformer CP colored in the circular permutation AFL, with the other CP in gray. **d**, Some permutation-tolerant residues (S52, N55, K57) in B-conformation CP mediate the interaction with RNA (tan). (PDB ID: 1ZDI)

According to the circular permutation AFL, permutants starting from the 52nd to the 58th residues in the MS2 CP sequence (referred to as CPM52 to CPM58, with the same nomenclature applying to other permutants as well) showed high AFSs that were higher than the CP-CP AFS, and therefore were identified as assembly-competent variants. CPM2 and CPM24 showed positive AFSs but at a lower value than the CP-CP AFS, indicating their slight enrichment after the size selection, which indicated an intermediate level of assembly competency. There were also some permutants with slightly negative AFSs (−1.0–0), including CPM29, CPM68, and CPM129, suggesting their moderate presence in the screened library. Interestingly, even though other permutants had extremely low AFSs, none of them were completely undetectable after the size selection (Supplementary Data). This suggests that none of the permutants lose their capability for VLP assembly entirely, but they differ in their overall competency to readily assemble into VLPs. We classified all permutants with negative AFS as assembly-deficient variants due to their reduction in relative abundance after the size selection.

To confirm that our circular permutation AFL provides insight into the assembly competency of every permutant, we randomly selected 13 variants and tested their assembly competency through individual SEC-based assays. The selected variants cover a wide range of AFSs and have new termini located at various secondary structure elements of MS2 CP (Fig. 3b). For example, permutants with high AFSs included CPM53, CPM56, and CPM57 from the super tolerant region, while CPM2 has an AFS of 0.22 (which is between 0 and the AFS for CP-CP, 0.48). CPM29 and CPM129 represent variants with slightly negative AFSs. Finally, the remaining seven variants are low-AFS permutants that contain new terminal locations covering the exterior βB strand, interior βF and βG strands, the FG-loop, the loop connecting βG strand and α1 helix, as well as the loop between α1 and α2 helices. This randomly selected subset of variants included both assembly-competent and assembly-deficient variants to validate our AFL calculation.

We quantified the assembly competency of selected permutants by calculating the VLP yield, as assessed by measuring the A_280_ peak area from the SEC chromatograph, for every selected variant purification (Supplementary Fig. 3). To ensure that our assembly assay calculation distinguishes assembly-deficient and assembly-competent variants (Fig. 3b), we used TEM to examine the compositions of purified particles from the defined MS2 VLP peak. Indeed, assembly-competent variants showed abundant VLP particles under TEM (Supplementary Fig. 4), indicating their strong propensity to self-assemble. Interestingly, we also found rare VLP particles from SEC-purified assembly-deficient variants (Supplementary Fig. 4). When we compared variant AFSs to their individually characterized assembly competency, all 13 variants exhibited consistent trends (Fig. 3b). For example, CPM2 did not exhibit substantial assembly, as expected for a variant with an intermediate-level AFS; variants with similar scores in previous MS2 CP library studies also exhibited low VLP titers^5,9,10,29^. Our observations suggest that permutants with high AFSs should assemble into VLPs easily and have great potential for further engineering and modification.

### Biophysical implications of circular permutation AFL

Secondary structures in the MS2 CP exhibit differences in their tolerance to circular permutation, which introduces new termini within the secondary structures (Fig. 3c). We hypothesized that disruptions to certain structure elements via introducing new termini would disrupt protein folding or assembly and lead to the apparent VLP assembly deficiency. In general, interior secondary structures were more tolerant to introducing new termini than exterior ones. This is likely due to the robustness of β sheets at the VLP interior, which are mostly stabilized by the hydrogen bonding network across neighboring strands. However, the exterior secondary structures were mostly short β strands and α helices whose stability could be easily disrupted by introducing the new termini in these locations. Meanwhile, structure elements at the protein-protein interfaces were less tolerant to hold new termini. Even though the FG-loop exhibited a high mutability in a previous mutagenesis study^9^ and showed tolerance to some three-residue peptide insertions^10^, permuted CP-CP constructs starting at the FG-loop failed to yield assembled VLPs. Similarly, other secondary structures engaged in protein-protein interfaces generally had low tolerance to new termini (Supplementary Fig. 5). On the contrary, the EF-loop was highly mutable per prior studies and is also quite tolerant to the new termini. Multiple residues in this region are involved in protein-nucleic acid interactions (Fig. 3c-d)^6,22,36,37^. This is surprising because such interactions are thought to be critical for assembly^21^ and encapsulation of cargo^28^. This result suggests that the interaction between the protein shell and charged cargo is driven by non-specific electrostatic interactions in the permutants.

When we compared our circular permutation AFL to the mutability index matrix in the previous site saturation mutagenesis study^9^, we found that the region that highly tolerates circular permutation was also highly mutable (Supplementary Fig. 6a-b). This indicates that those permutants should have high flexibility for modification and engineering. On the contrary, not all highly mutable sites exhibited tolerance to circular permutation. However, after filtering out permutants with AFS ≤ -0.6, we observed a correlation (r = 0.79) between the AFS of the permutant and the mutability index of the permutation starting residue (Supplementary Fig. 6c). In our circular permutation library, AFS = -0.6 guaranteed the moderate presence of a permutant with at least 10^3^ read counts and at most 3/4 reduction after the size selection, where the AFS quantification should be reliable. This filtered correlation probably reflects the fact that our circular permutation AFS could be decomposed into two factors – the capability of the permutant to fold properly into the CP dimer conformation and the assembly favorability of the appropriately folded CP dimer. One explanation for this result is that for AFS ≤ -0.6, the low AFS may reflect a CP folding problem, while an AFS above -0.6 mainly reveals the favorability of the folded permutant towards VLP assembly.

### Structure characterization of the assembly-competent permutant

To confirm that the assembly-competent permutants assemble into VLPs in a similar pattern to the WT MS2 VLP, we solved the structure of the assembled CPM58 VLP at 2.6 Å using cryogenic electron microscopy (cryo-EM) (Supplementary Fig. 7 and Table 2). Similar to the WT MS2 VLP, circularly permuted CP-CP adopted two different conformations, with the asymmetric subunits mediating the icosahedral five-fold axes contacts on the pentagonal facets, and symmetric subunits interdigitating with asymmetric subunits around the icosahedral three-fold axes on the hexagonal facets (Fig. 4a-b). The symmetric subunit contains two symmetric C-conformer CP folds, and the asymmetric subunits comprise A- and B-conformer CP folds. CPM58 subunit models exhibit a high structural similarity to CP models in the WT MS2 VLP, with major differences at the orientation of the original CP N-terminus due to the designed fusion (Fig. 4c).

**Figure 4.**
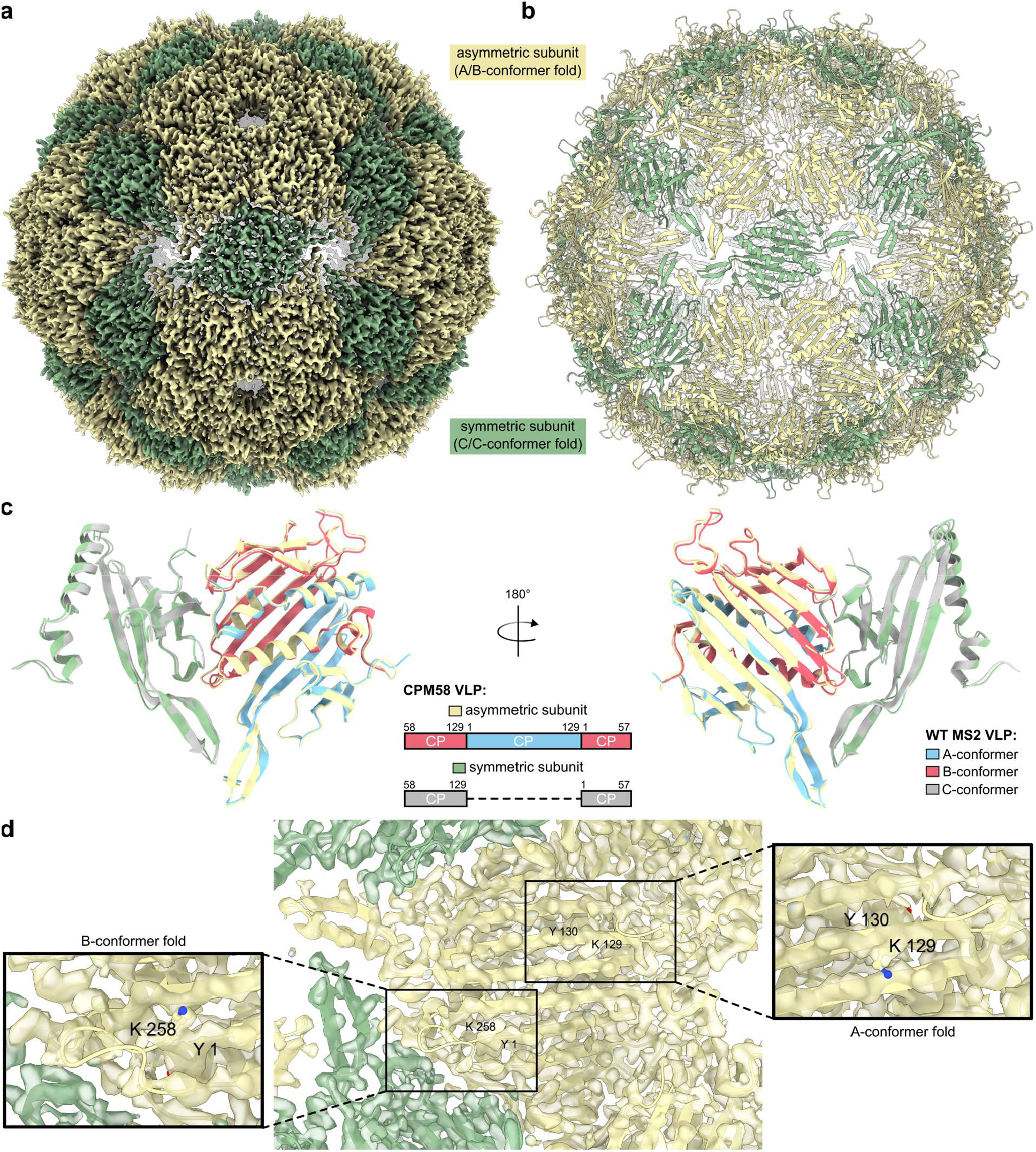
Cryo-EM structure of the assembled CPM58 VLP. **a,b**, Cryo-EM density map (**a**) and model (**b**) of the assembled CPM58 VLP with asymmetric and symmetric subunits colored diferently. **c**, CPM58 subunit models aligned to WT MS2 CP models (PDB ID: 1ZDI) with conformer folds in the WT models colored diferently. Residues are numbered as in the WT CP sequence. **d**, Zoomed-in view of the cryo-EM density map around the asymmetric subunit model. Residues corresponding to K57 and Y58 in the WT CP sequence are labeled and highlighted in ball-stick.

Circularly permuting CP-CP introduces sequence heterogeneity to the dimeric CP fold subunit -one CP fold in the subunit holds the new N- and C-termini with a structural gap in between, while the connectivity of structure elements of the other CP fold stays uninterrupted. Interestingly, we observed symmetric missing density at the designed terminal region of both CP folds in the symmetric subunit (Supplementary Fig. 8a), suggesting that the sequence heterogeneity has little effect on the symmetric subunit conformation. Because the two CP folds within the symmetric subunit are almost identical, we modeled the two C-conformer CP folds as separate single chains and utilized the VLP symmetry to build the whole assembly model (Fig 4c). However, in the asymmetric conformer we observed different density map continuity at the designed terminal regions in A- and B-conformer folds (Fig. 4d). There is a clear density break in the B-conformer CP fold, which resembles the map pattern in the C-conformer fold. On the contrary, the density map of the βF strand in the A-conformer fold is continuous, indicating a relatively fixed conformation of the intact βF strand. The difference in the βF strand density map continuity suggests a higher flexibility of the EF-loop structure in the B-conformer fold. We then reasoned that it was more likely for the B-conformer fold to hold the new termini of CPM58 while the A-conformer fold adopts the intact CP sequence. Therefore, we modeled the asymmetric subunit as a whole chain with termini positioned at the favored B-conformer fold. Apart from the EF-loop, AB-loops in all CP folds exhibited high flexibility with high B-factors and missing density in the cryo-EM map (Supplementary Fig. 8b-d), supporting the possibility for peptide fusion and insertion at those highly flexible regions.

### Assembly-competent permutants enabling terminal tagging

Given the high terminal flexibility of CPM58 in the cryo-EM structure, we hypothesized that the assembly-competent permutants would be tolerant of terminal extension and peptide display. To test our hypothesis, we fused the 13-amino acid SpyTag to the C-termini of top-ranked permutants, including CPM2 and CPM52–58, via -GSS-linkers (Fig. 5a)^38,39^. We then characterized VLP particles with SEC, SDS-PAGE, negative stain TEM, and DLS. With the exception of CPM2 and CPM55, the selected permutant-SpyTag constructs exhibited high assembly competency with successful incorporations of SpyTag at the VLP interior (Fig. 5b-c). The sizes of assembled permutant-SpyTag VLPs showed similar diameters to WT MS2 VLPs, as measured by DLS (Supplementary Table 1). While the majority of VLPs had normal sizes, we observed small proportions of larger VLPs and stretched VLPs via TEM (Fig. 5d). Those abnormal VLPs could be in T = 4 or other non-icosahedral, lower-symmetry structures, indicating that the presence of C-terminal SpyTag may have a mild effect on slowing down the transition from C/C to A/B conformation and therefore affect the VLP self-assembly^39^. In addition, permutant-SpyTag VLPs seem to maintain the ability of nucleic acid encapsulation with high absorbance at 260 nm (Fig. 5c), suggesting it would be possible to use a high-throughput setup to screen for proteins or peptides that can be fused to the assembly-competent CP-CP permutants. Since the WT MS2 CP is intolerant to SpyTag terminal extension (Supplementary Fig. 9), our results indicate that circularly permuting the MS2 CP-CP construct enables terminal tagging and introduces novel structures for MS2-based VLPs.

**Figure 5.**
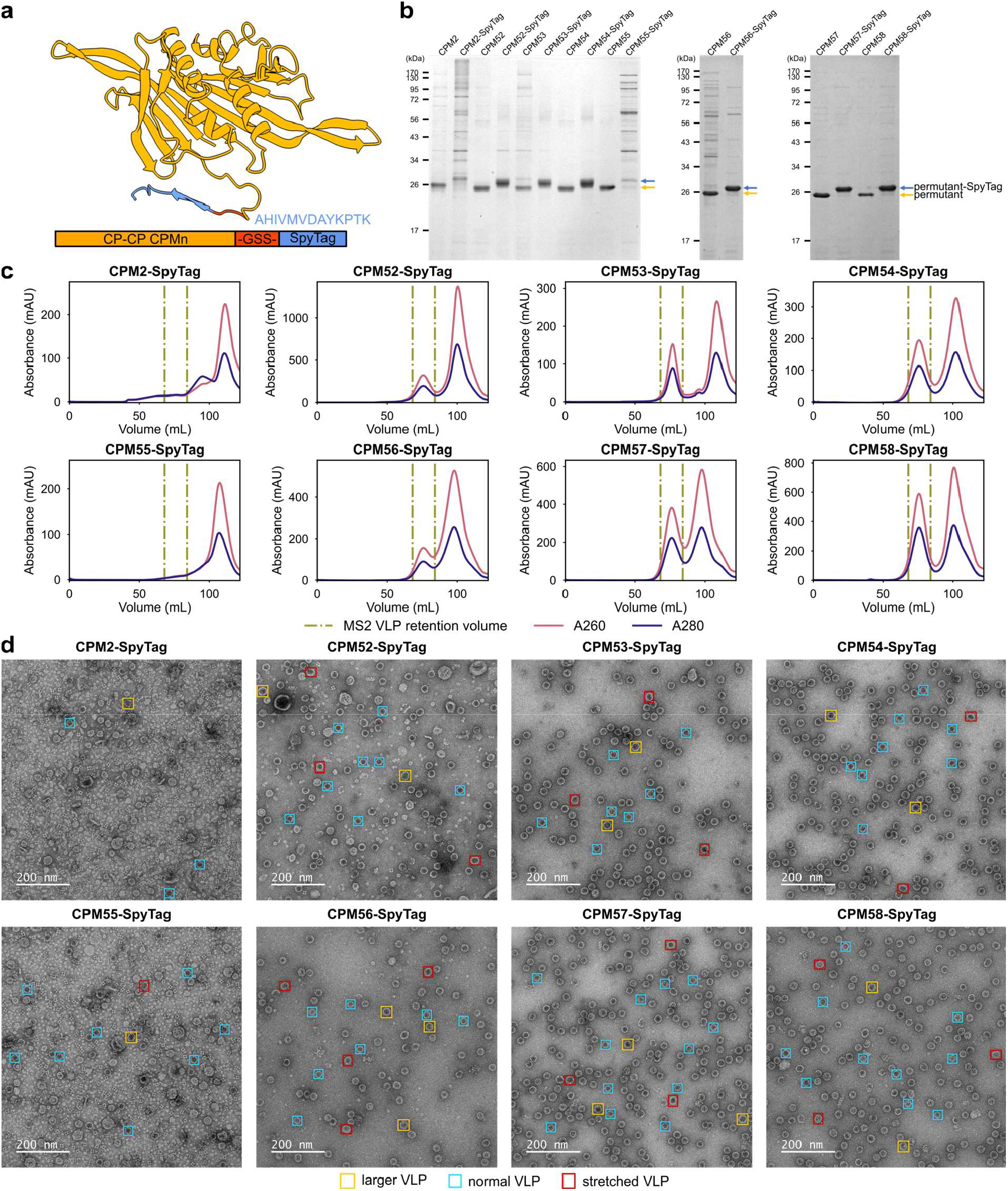
C-terminal SpyTag fusion on MS2 permutants. **a**, Design of permutant-SpyTag constructs. The 13-amino acid SpyTag was fused to the C-terminus of the permutant via a -GSS-linker. AlphaFold3 predicted CPM58-SpyTag is shown. **b**, SDS-PAGE gel of purified permutant-SpyTag VLPs in comparison to purified permutant VLPs. **c**, SEC chromatographs of permutant-SpyTag VLP purifications. **d**, TEM micrographs of purified permutant-SpyTag VLPs

### A new protein encapsulation strategy with permutants

We hypothesized that the C-terminal SpyTag would maintain its reactivity in forming the covalent isopeptide bond with SpyCatcher^38,39^. Therefore, we co-expressed a SpyCatcher-sfGFP protein with assembly-competent permutant-SpyTag fusion, purified assembled VLPs with SEC, and characterized them with SDS-PAGE, negative stain TEM and DLS (Fig. 6a). With the exception of CPM56-SpyTag/SpyCatcher-sfGFP, all co-expressions showed substantial VLP peaks on SEC, with a high absorbance at 260 nm and an accumulation of green fluorescence, indicating the presence of nucleic acids as well as the green-fluorescent cargo (Fig. 6c). SDS-PAGE of purified VLPs confirmed the covalent bonding between SpyTag and SpyCatcher as well as the presence of excess permutant-SpyTag (Fig. 6b). We observed relatively homogenous VLPs with the majority being T = 3 VLPs with normal sizes, as assessed by DLS and TEM (Fig. 6d and Supplementary Table 1). Taken together, our results suggest that we can encapsulate protein cargo inside MS2 VLPs via the SpyTag/SpyCatcher interaction along with its intrinsic nucleic acid encapsulation capability, and that the presence of protein cargo may only have a mild effect on the morphology of assembled VLPs.

**Figure 6.**
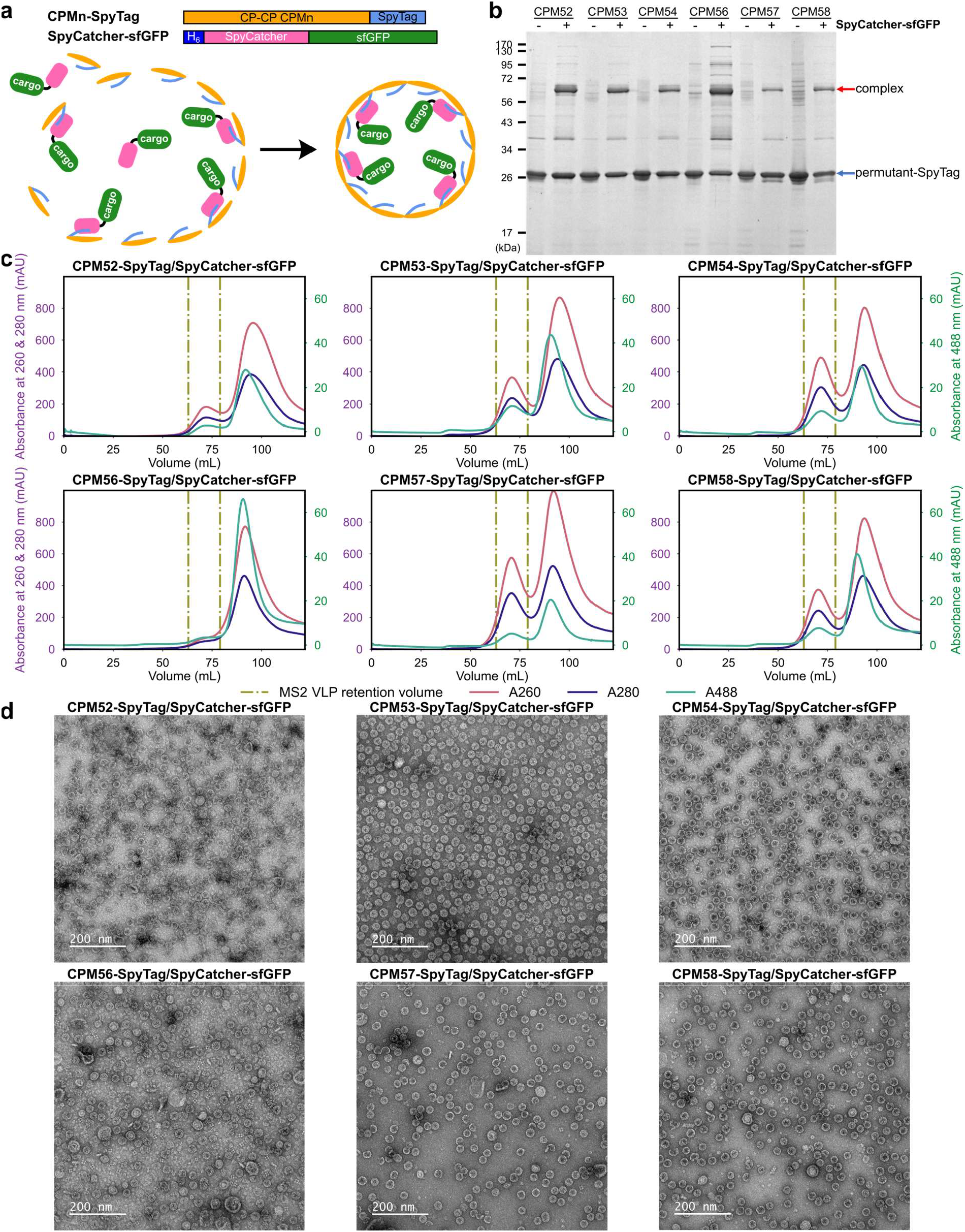
A protein encapsulation system with MS2 permutants. **a**, Diagram of in vivo cargo (sfGFP) encapsulation via co-expression. **b**, SDS-PAGE gel of purified permutant-SpyTag/SpyCatcher-sfGFP VLPs in comparison to purified permutant-SpyTag VLPs. The covalently bound permutant-SpyComplex-sfGFP band is labeled as complex. **c**, SEC chromatographs of permutant-SpyTag/SpyCatcher-sfGFP VLP purifications. **d**, TEM micrographs of purified permutant-SpyTag/SpyCatcher-sfGFP VLPs.

## Discussion

Systematic engineering of protein assemblies through genetic modifications is powerful in revealing assembly principles and guiding engineering for myriad applications. In nature, circular permutation is a widely used strategy to diversify the protein sequence space and potentially adopt novel protein functions. However, this strategy has not been exploited in the test tube as frequently as other protein engineering strategies such as substitution, deletion and insertions^8^. Here, we performed a comprehensive circular permutation study on a self-assembling protein system for the first time. We combined a systematically generated circular permutation library on MS2 CP-CP with size selection for VLPs to explore the full sequence and structure space and identify positions that tolerate introducing new termini. Compared to the site mutagenesis and short loop insertion libraries on VLPs, circularly permuting CP-CP applies more stress on the CP dimer folding but still reflects VLP assembly competency from properly folded subunits^4,5,9,10,29^. Indeed, our circular permutation AFL showed different tolerance patterns from previous MS2 studies. For example, the highly mutable FG-loop exhibited low tolerance towards relocating termini. We suggest that this is due to the folding pitfall of permutants starting at the flexible FG-loop region, which emphasized the essential role of the appropriate protein folding prior to the VLP assembly^9,10^. Moreover, our cryo-EM structure of the assembly-competent CPM58 VLP revealed a similar subunit arrangement pattern to the WT MS2 VLP, indicating that overcoming the folding barrier is the main obstacle for a circularly permuted construct to form VLPs. This underscores the fact that this circular permutation study can provide novel insights into MS2 VLP assembly. Meanwhile, our systematic characterization identified novel positions from past peptide insertion studies – the EF-loop and the beginning of βF strand – exhibited great potential in placing new termini, providing an alternative to the popular insertion loci at AB- and FG-loops^10,34,39–42^. Given that the EF-loop in the WT MS2 CP can tolerate the SpyTag insertion with a carefully designed linker^33^, we believe our circular permutation study provides guidance for further peptide insertions on the MS2 CP.

At the same time, we have demonstrated the great potential of circular permutation in diversifying the structure space of protein subunits and engineering a proteinaceous nanoparticle for novel functions. Genetic peptide fusion or insertion has been difficult for the MS2 CP, as such modifications usually come with failures in VLP assembly or changes in the VLP morphology^5,33^. However, we were able to easily fuse the SpyTag to C-termini of most assembly-competent permutants and maintain a relatively homogenous VLP population. In addition, we observed missing density around the termini of CPM58 in the cryo-EM map of the assembled VLP, suggesting the high flexibility of the new termini in their orientations. On that account, the fused tag to our circularly permuted constructs may adopt a favored spatial pattern inside the VLP, as CPM58 exhibited a preference for positioning the terminal opening in the B-conformer CP fold of the asymmetric subunit. The successful C-terminal SpyTag extension on CPM53 is consistent with a previous SpyTag insertion study on the monomeric MS2 CP^33^. Unlike results from inserting SpyTag into the monomeric CP^33^, our CPM53-SpyTag can assemble into nanoparticles without the unexpected rod population, perhaps a benefit from having only one SpyTag copy per CP dimer fold in the CPM53-SpyTag VLP. Taken together, our assembly-competent permutants offer a promising solution for terminal tagging and even protein fusion.

Moreover, the attached SpyTag exhibited reactivity to covalently bond to SpyCatcher, which indicates a high flexibility of the terminal tag. This also provides a new protein encapsulation strategy with a broad spectrum of protein cargos utilizing the SpyTag/SpyCatcher system^38^, as there is no requirement for introducing negative charges to cargo surface. This could be utilized in various applications, such as optical imaging^43^, antigen encapsulation^44^, and enzymatic pathway compartmentalization^33^. Finally, the assembled permutant VLPs maintain the ability to encapsulate nucleic acids, enabling possible multivalent cargo encapsulations by combining nucleic acid tagging on the cargo and terminal modifications on MS2 CPs.

More work will be needed to uncover design principles of self-assembling proteinaceous nanoparticles and further expand the sequence and structure space of engineered proteinaceous nanoparticles. Specifically, it is attractive to consider combining circularly permuted CP-CP with mutagenesis. Hilvert and coworkers evolved a protein cage made of circularly permuted protein subunits into an artificial nucleocapsid via directed evolution^19^, raising the possibility that our assembly-competent circular permutants may also provide promising starting platforms to tailor protein sequence and structure with desired function. This in turn would help establish more comprehensive VLP assembly rules, such as studying epistatic interactions across monomeric CP folds. It is possible that point mutations across the MS2 CP could stabilize circular permutants with moderate assembly competency, and vice versa. Circularly permuted CP-CP might also expand the engineering capacity of MS2 VLPs through orthogonal modifications, for example, coupling terminal tagging with mutations and chemical conjugations. Further combinational modifications on MS2 VLPs and other proteinaceous nanoparticles will expand the potential of protein-aided nanotechnology.

## Methods

### Strains and Plasmids

Chemically competent *E. coli* DH10B cells and the pBAD33t vector were used for the unpermuted CP-CP construct, library construction, and expression as well as individual assembly assays for library validation. Chemically competent *E. coli* BL21 (DE3) cells were used for the expression of assembly-competent permutants and permutant-SpyTag constructs, as well as the co-expression of permutant-SpyTag and SpyCatcher-sfGFP. The pET28b(+) vector was used for cloning assembly-competent permutants and permutant-SpyTag constructs, and the pET21b(+) vector was used for cloning the SpyCatcher-sfGFP construct. All vectors contained BsaI cut sites for Golden Gate (GG) Assembly to insert designed constructs. Detailed information about the strains and plasmids can be found in the Supplementary Data.

### VLP Expression and Purification

To express the CP-CP construct, the whole circular permutation library, and the selected permutants for validation, glycerol stocks were inoculated into LB broth, Miller (Fisher Scientific, Cat# BP1426-500), containing 34 μg/mL chloramphenicol (1 L for the pooled library expression and 50 mL for individual constructs) and allowed to grow to an OD_600_ of 0.6–0.8 at 37°C with shaking. The culture was then cooled down to 16°C and the protein expression was induced by arabinose at a working concentration of 0.02% (w/v). Expression proceeded at 16°C with shaking overnight and cells were then harvested and sonicated. Expressed protein particles went through an ammonium sulfate precipitation at half saturation. Precipitates were re-suspended in size exclusion chromatography (SEC) buffer (200 mM sodium chloride, 20 mM sodium phosphate, pH 7.2) and dialyzed against the same buffer overnight. The soluble portion was purified in the SEC buffer by HiPrep 16/60 Sephacryl S-500 HR column (Cytiva, Cat# 28935606) on an Akta Pure 25 L fast protein liquid chromatography (FPLC) system. Fractions within the MS2 VLP retention volume were collected, combined, concentrated using 100kDa Vivaspin columns (Fisher Scientific, Cat#28-9323-63), and subjected to SDS-PAGE, DLS, and TEM analyses.

For the expression of assembly-competent permutants and permutant-SpyTag constructs, we carried out a 50 mL expression in LB broth, Miller, with 50 μg/mL kanamycin at 37°C with shaking. Cultures were induced when the OD_600_ reached 0.6-0.8 using 400 μM isopropyl β-D-1-thiogalactopyranoside (IPTG) induction, and from this point onward protein expression and VLP purification proceeded as described above for the library. Similarly, to co-express permutant-SpyTag and SpyCatcher-sfGFP, corresponding plasmids were co-transformed into *E. coli* BL21 (DE3). Cells were plated on LB agar plates with 50 μg/mL kanamycin and 50 μg/mL carbenicillin for selection. A 50 mL protein expression with dual antibiotics and IPTG induction as well as VLP purification in the SEC buffer were then carried out as described above.

### Negative stain transmission electron microscopy (TEM)

Samples were prepared for TEM analysis by diluting to approximately 0.1 mg/mL protein prior to the grid preparation. 10 μL analyte solutions were then applied to glow discharged carbon-coated copper grids (Electron Microscopy Sciences, Cat# FCF400-Cu-50) for about 10 s and wicked away. The grids were then exposed to 10 μL negative stain solution (1% uranyl acetate) twice, which was immediately wicked away. Another 10 μL negative stain solution was then added onto the grids for 4 min and wicked away completely. Images were obtained using the JEOL 1400 Flash TEM with a Gatan OneView Camera at 120 kV accelerating voltage.

### Sodium Dodecyl Sulfate-polyacrylamide Gel Electrophoresis (SDS-PAGE)

We made 12.5% SDS-PAGE gels and loaded samples that had been pre-mixed with Laemmli buffer. Gels were run with SDS-PAGE running buffer (3 g/L Tris, 14.4 g/L glycine, 1 g/L SDS) for 80 min at 160 V, stained with the Coomassie Blue stain, and imaged with the Azure 600 Imager.

### Dynamic Light Scattering (DLS)

Samples were prepared by diluting to approximately 0.1 mg/mL protein prior to the DLS measurement. Particle sizes were then analyzed using Malvern Zetasizer Ultra (light source: He-Ne laser 633 nm, nominal power: 4 mW) with a protocol for proteins in phosphate-buffered saline (PBS) at 25°C. Each size measurement contained 15 size runs with durations of 6.71s, and the fluorescence filter was used to eliminate interference by possible fluorescent cargos. For every sample, 3 successful size measurements were used to determine the particle size and standard deviation. Reported values were diameters of peak 1 ordered by area in the volume distribution.

### Library Generation

The 129-variant circular permutation library was divided into 5 sub-libraries (CPM2–27, CPM28– 53, CPM54–79, CPM80–105, and CPM106–129 + CP-CP) and unique 4-base overhangs were designed to direct the assembly of every CP-CP permutants within one sub-library. Using the CP-CP sequence as a template, individual polymerase chain reactions (PCRs) were used to obtain CP encoding fragments with designed GG overhangs (Supplementary Data). Amplified, double-stranded CP permutant DNA copies were subjected to DpnI (NEB, Cat# R0176S) digestion and purified using a PCR Clean-up Kit (Promega, Cat# A9285). Five pooled GG cloning reactions were constructed to assemble all permutants in a sub-library at once. GG products from 5 sub-libraries were then combined, transformed into chemically competent DH10B cells and then plated on large (150×15 mm, Fisher, Cat# 08757148) LB agar plates with 34 μg/mL chloramphenicol. The number of colonies varied among biological triplicates, but every transformation yielded a at least 10 times more colonies than the theoretical library size. The correct assembly of CP-CP permutants was confirmed by picking up colonies randomly followed by Sanger sequencing. Colonies were then scraped from the entire plates, transferred into LB broth, Miller, allowed to grow for at least 5 h, and stored with glycerol at -80°C.

### Sample Preparation for High-throughput Sequencing

For the unscreened library, plasmid DNA was extracted from a 5 mL overnight library culture with the Zyppy Plasmid Miniprep Kit (Zymo Research, Cat# D4037). For the screened library, RNA was extracted from the purified VLPs as previously published protocols^5,9,10,29,45^. An equivalent volume of TRIzol (Invitrogen, Cat# 15596018) was added to the concentrated purified VLPs and subjected to heat for homogenization. Then, chloroform was added to the supernatant. After a centrifugation at 16,000 ×g for 15 min, the liquid was separated to 3 phases. The RNA in the upper aqueous phase was collected, precipitated by adding isopropanol, washed with 70% ethanol, and finally re-suspended in RNase-free water. cDNA was synthesized from the extracted RNA with SuperScript™ III First-Strand Synthesis System (ThermoFisher, Cat# 18080051). DNA samples from screened and unscreened libraries were then amplified with two rounds of PCR to add barcodes (10 cycles) and the Illumina sequencing handles (6 cycles). Samples were combined and analyzed by Miseq Reagent Kit v3 (600 cycle) and an overall Q30 of about 42% was obtained.

### High-throughput Sequencing Data Processing

Data was trimmed using Trimmomatic^46^ with a 6-unit sliding quality window of 15 and a minimum length of 30. To align reads to every possible permutants, a reference library containing all circularly permuted CP-CP sequences with short upstream and downstream plasmid sequences was prepared, where every reference sequence was named after the starting residue number of the corresponding permutant. Forward reads from the filtered data were then extracted and aligned to the reference library with Burrows-Wheeler Aligner (BWA-MEM)^47^. Reads were then sorted and indexed with Samtools^48^, and unmapped reads were filtered out with the Picard function CleanSam. The identity of every aligned read was then confirmed by comparing the first 8-nt sequence downstream of the start codon to the reference sequence it aligned to, using Python scripts written in house.

### Apparent Fitness Landscape (AFL) Calculation

The calculation of AFL was similar to protocols described in previous publications^5,9,10,29^. Percent abundance within a library was calculated as the ratio of a count for a specific permutant to all permutant counts in the library. Relative percent abundances were then calculated by dividing percent abundances of selected libraries by corresponding percent abundances of unscreened plasmid libraries. Rare relative percent abundances were replaced with 0.0001 to enable the following calculation. Log10 of the relative percent abundance was then defined as the AFS. AFS matrices were calculated from three biological replicates, and the final AFS in the AFL was calculated as the average AFS across three replicates. Standard deviations of AFS were also calculated across triplicates.

### Individual Assembly Competency Assay

The assembly competency assays were done in biological triplicates, and the values reported were the average VLP peak area with standard deviations as error bars. SEC fractions at VLP peaks were analyzed with SDS-PAGE and the purest fractions were subjected to TEM sample preparations.

### Cryo-electron Microscopy (cryo-EM)

#### VLP expression and purification

CPM58 was cloned onto pET28b(+) vectors, transformed into *E. coli* BL21 (DE3) cells for protein expression. Proteins were then expressed in LB broth, Miller, with 50 μg/mL kanamycin and cells were allowed to grow to an OD of 0.6–0.8 at 37°C. The culture was then cooled down to 16°C and the protein expression was induced by adding 400 μM IPTG. Expression proceeded at 16°C overnight and cells were then harvested and sonicated. VLPs were purified similarly to described above except that the SEC buffer was replaced with a buffer composed of 100mM sodium chloride, 20 mM HEPES, pH 7.2. Purified VLP fractions were collected, spin-concentrated, and frozen with liquid nitrogen until sample preparation.

#### Sample preparation

The purified VLPs were concentrated to 2.0 mg/mL. Quantifoil R1.2/1.3 400 mesh holey carbon grids (Quantifoil Micro Tools) were glow-discharged using a Pelco easiGlow glow discharge system (Ted Pella) at 15 mA for 30s. Aliquots (3 μL) of protein samples were applied to the grids and blotted at 15°C, 95% relative humidity level and 0 blot force for 4s, using a Vitrobot Mark IV blotting system (ThermoFisher Scientific) before plunging in liquid ethane for vitrification.

#### Data collection

Movies of MS2 VLP particles embedded in vitreous solution were collected at liquid nitrogen temperature using Glacios transmission electron microscopes (ThermoFisher Scientific). The microscope was equipped with a High Brightness Field Emission Gun (X-FEG, ThermoFisher Scientific), Selectris energy filter (ThermoFisher Scientific) operated at 10 eV energy spread width and a Falcon4i direct electron detector (ThermoFisher Scientific). The Falcon4i movies were recorded using EPU at 130,000X magnification with 0.894 Å/pixel pixel size. A defocus range of -0.6 – -1.6 μm was applied. The total electron dose of each movie was 60 e^-^/Å^2^. The data collection parameters are in Supplementary Table 2.

#### Data processing

Beam-induced motions of particles were corrected in patches using cryoSPARC v4^49^. Contrast transfer function (CTF) parameters were estimated using Gctf^49^. Particles were automatically picked *ab initio* using soft-edged disk templates internally generated on cryoSPARC v4^49^, followed by template picking using best 2D class averages. The picked particle images were boxed, extracted and 2D classified using cryoSPARC v4^49^. 3D classification without alignment was carried using cryoSPARC v4^49^. The initial 3D reconstructions were carried out ab initio, followed by non-uniform 3D refinement using cryoSPARC v4^49^ with refinement of per-particle defocus and per-group CTF parameters. The number of movies and particles used towards final reconstructions are in Supplementary Fig. 7.

A previous model of WT MS2 (PDB ID: 1ZDI)^37^ was used as an initial model and manually rebuilt using COOT^50,51^. All protein models were real-space refined using Phenix-1.21.2^52^, and evaluated using COOT and MolProbity server^53^. The cryo-EM maps were deposited in Electron Microscopy Databank (EMDB) and the coordinates of the atomic models were deposited in Protein Data Bank (PDB)^54,55^. The figures were generated using Chimera and ChimeraX^56,57^.

## Data Availability

The atomic coordinates and cryo-EM density maps will be available at the PDB and EMDB under accession codes PDB 9OH5 & EMDB-70484.

## Code Availability

Codes used for sequencing data processing and AFL calculation are available on GitHub (https://github.com/DTElab/MS2-circular-permutation).

## Acknowledgements

We thank all members of the Tullman-Ercek lab for their helpful discussions and support, especially Dr. Yu Liu for his advice on how to optimize many of the experiments. We thank Edric K. Choi and Jiayu Fu from the Lucks lab at Northwestern University for their assistance with MiSeq. We thank Dr. Pamela Focia and Prof. Alfonso Mondragón at Northwestern University Structural Biology Facilities (SBF) for helpful discussion and advice.

We gratefully acknowledge funding from the National Science Foundation grant CBET-2043973 to D.T.-E. S.L. was partially funded by the Molecular Biophysics Training Program grant awarded to Northwestern University (NIH 5T32GM140995-04). D.C.A. was funded by the Chemistry of Life Processes Predoctoral Training Program grant awarded to Northwestern University (NIH T32GM149439). N.W.K. was funded by a grant from the Department of Energy (DOE DE-SC0022180). This work used resources of the Northwestern University Structural Biology Facility, which is generously supported by NCI CCSG P30 CA060553 grant awarded to the Robert H. Lurie Comprehensive Cancer Center.

## Author contributions

S.L. and D.T.E. conceived this project. S.L., K.B., D.C.A., and J.J. performed experiments with results shown in this manuscript. S.L., N.W.K., and D.T.E. analyzed and interpreted experimental data. S.L. wrote the manuscript. S.L., N.W.K., and D.T.E. edited the manuscript. All authors reviewed and contributed to the manuscript.

## Competing Interests

The authors declare no competing interests.

## Notes

### Competing Interest Statement

The authors have declared no competing interest.

