## Supplementary Figures and Tables for "Synthetic Rewiring of Virus-Like Particles via Circular Permutation Enables Modular Peptide Display and Protein Encapsulation"

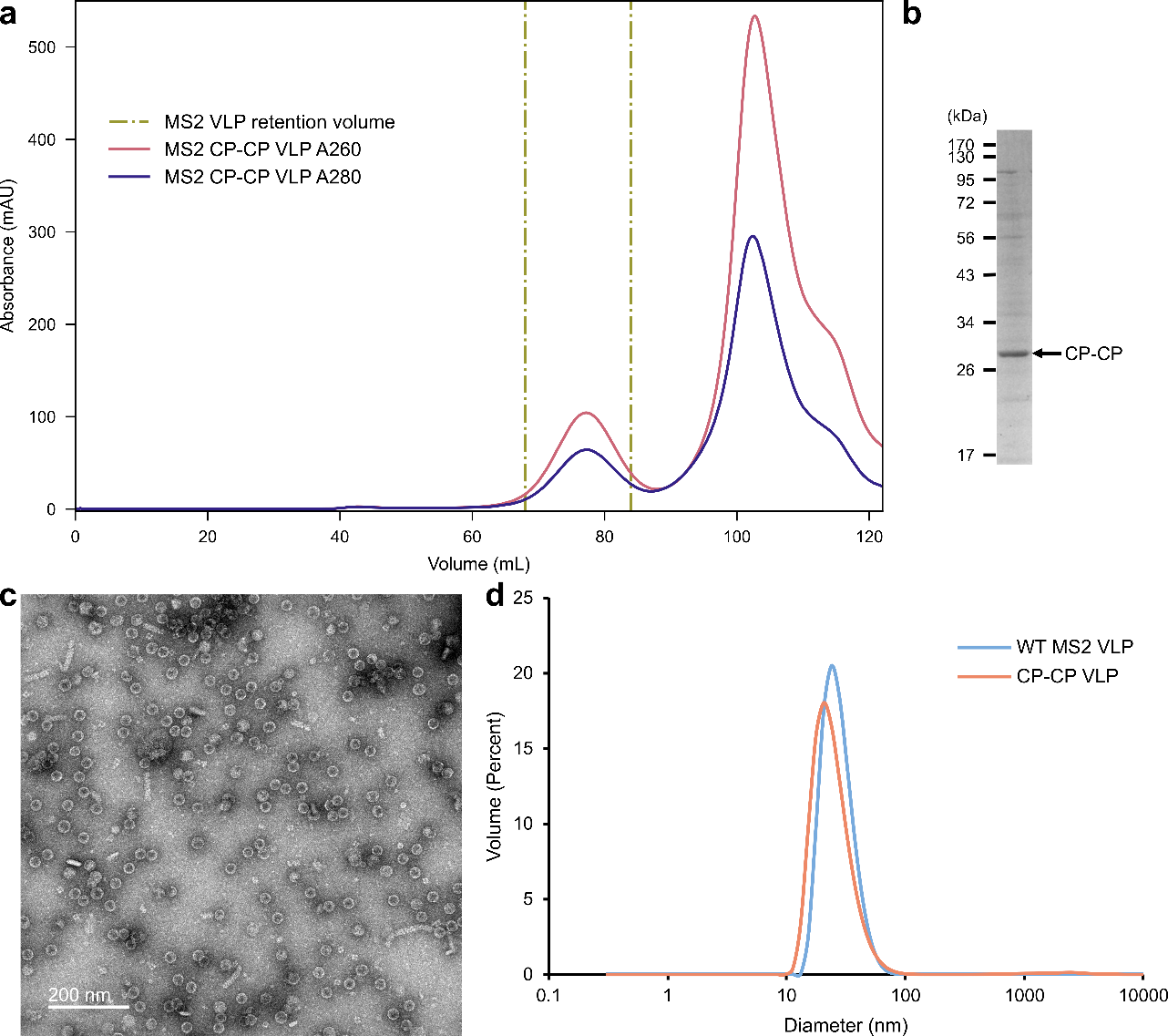

***Supplementary Fig. 1. Purification and characterization of CP-CP VLPs.*** **a**, The SEC chromatograph of the CP-CP VLP purification. **b-d**, SDS-PAGE (**b**), TEM micrograph (**c**), and DLS (**d**) of purified CP-CP VLPs.

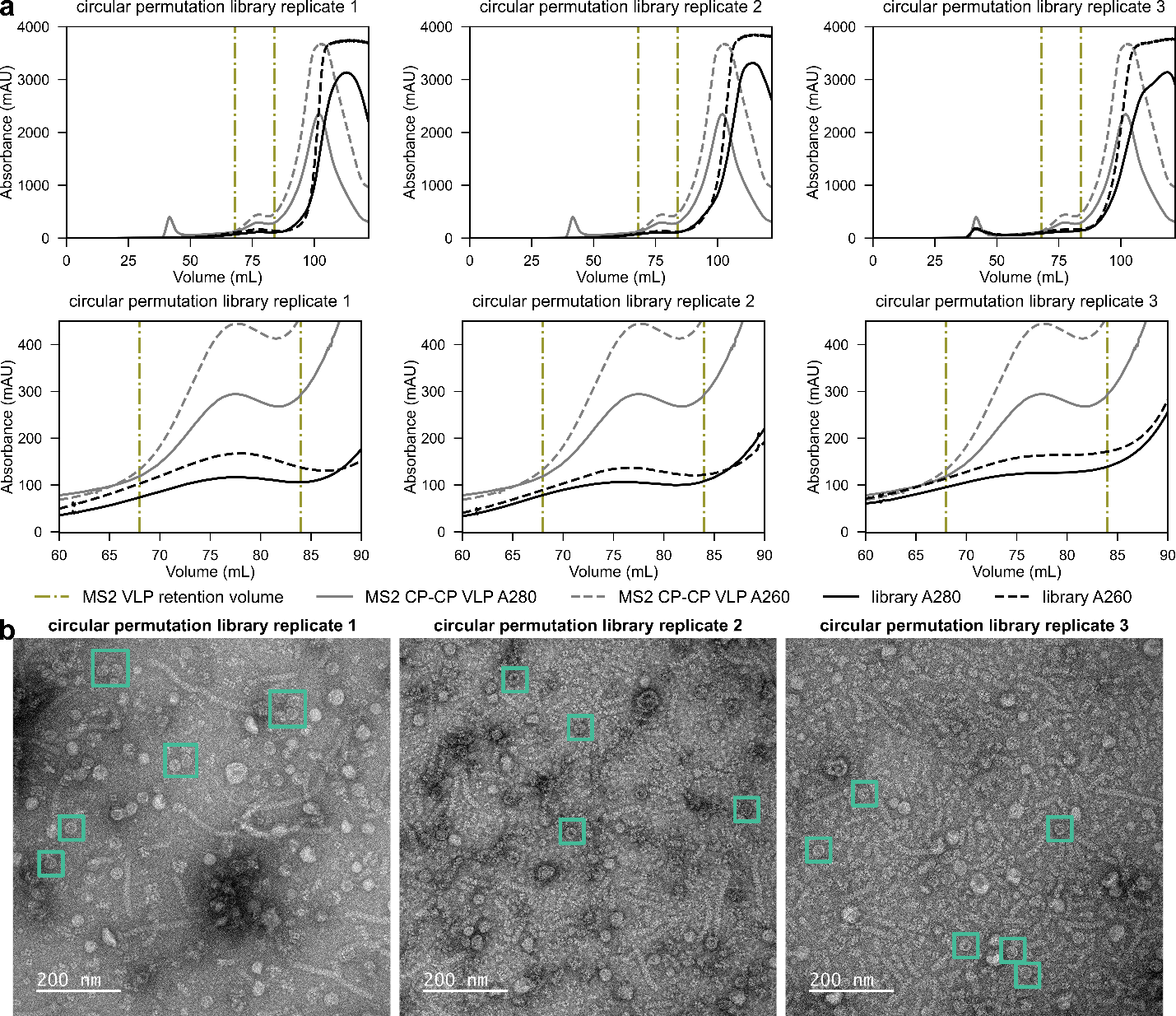

***Supplementary Fig. 2. SEC chromatographs and negative stain TEM of CPM library purification in biological triplicates.* a**, SEC chromatographs of three SyMAPS replicates with comparison to the purification of CP-CP under the same expression condition. Zoomed-in views of the VLP peak are shown under the whole chromatographs. Samples eluted between the two orange dotted lines were collected for high-throughput sequencing analysis. **b**, TEM micrographs of purified CPM libraries from SEC. Examples of VLP-like structures are boxed in mint. Samples were stained with uranyl acetate and imaged under the 50,000X magnification rate.

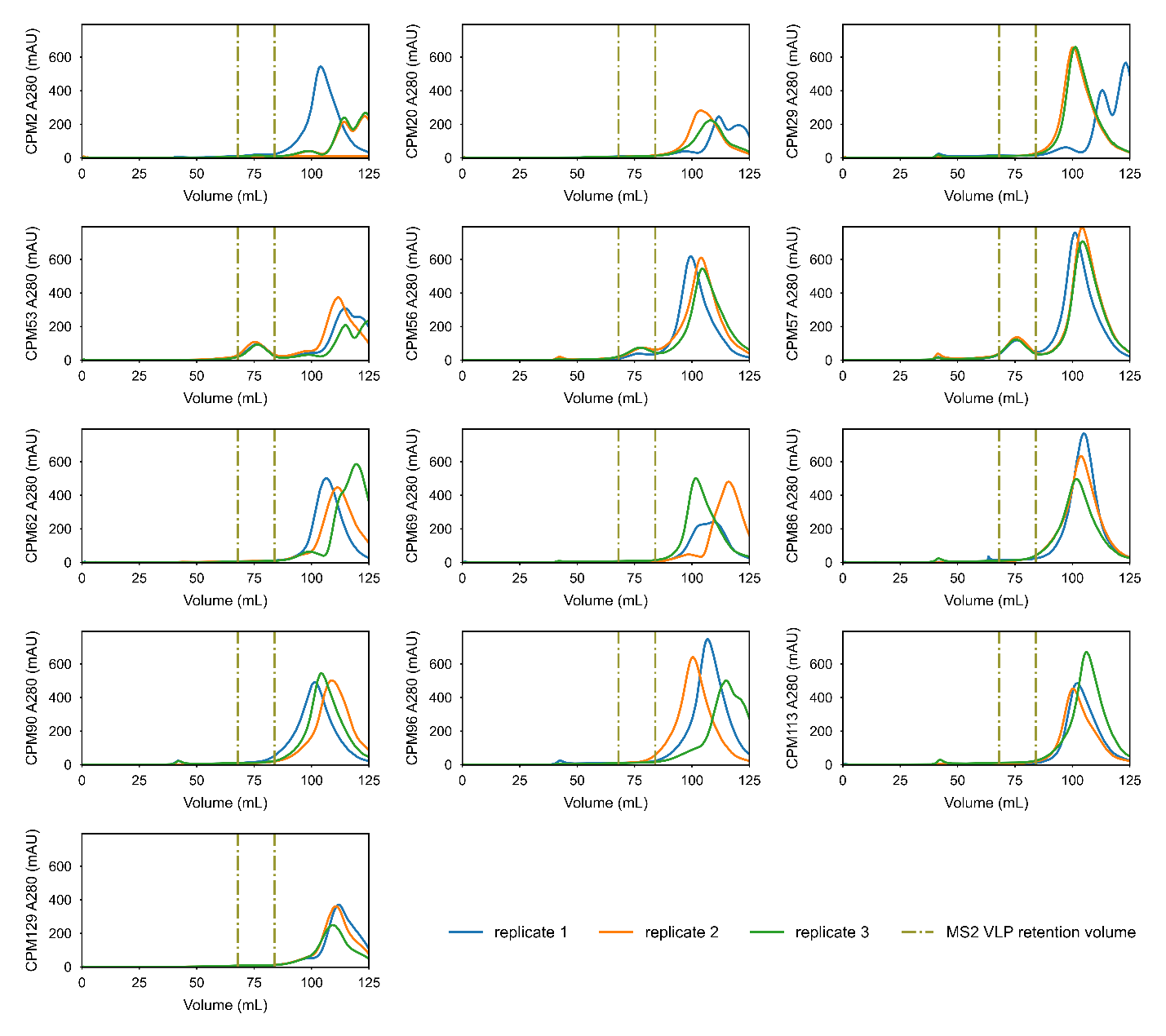

***Supplementary Fig. 3. SEC chromatographs of selected permutants for individual assembly assays.*** Chromatographs at A280 were shown in biological triplicates. Regions between the two red dotted lines were selected for VLP peak area calculation for assembly quantification.

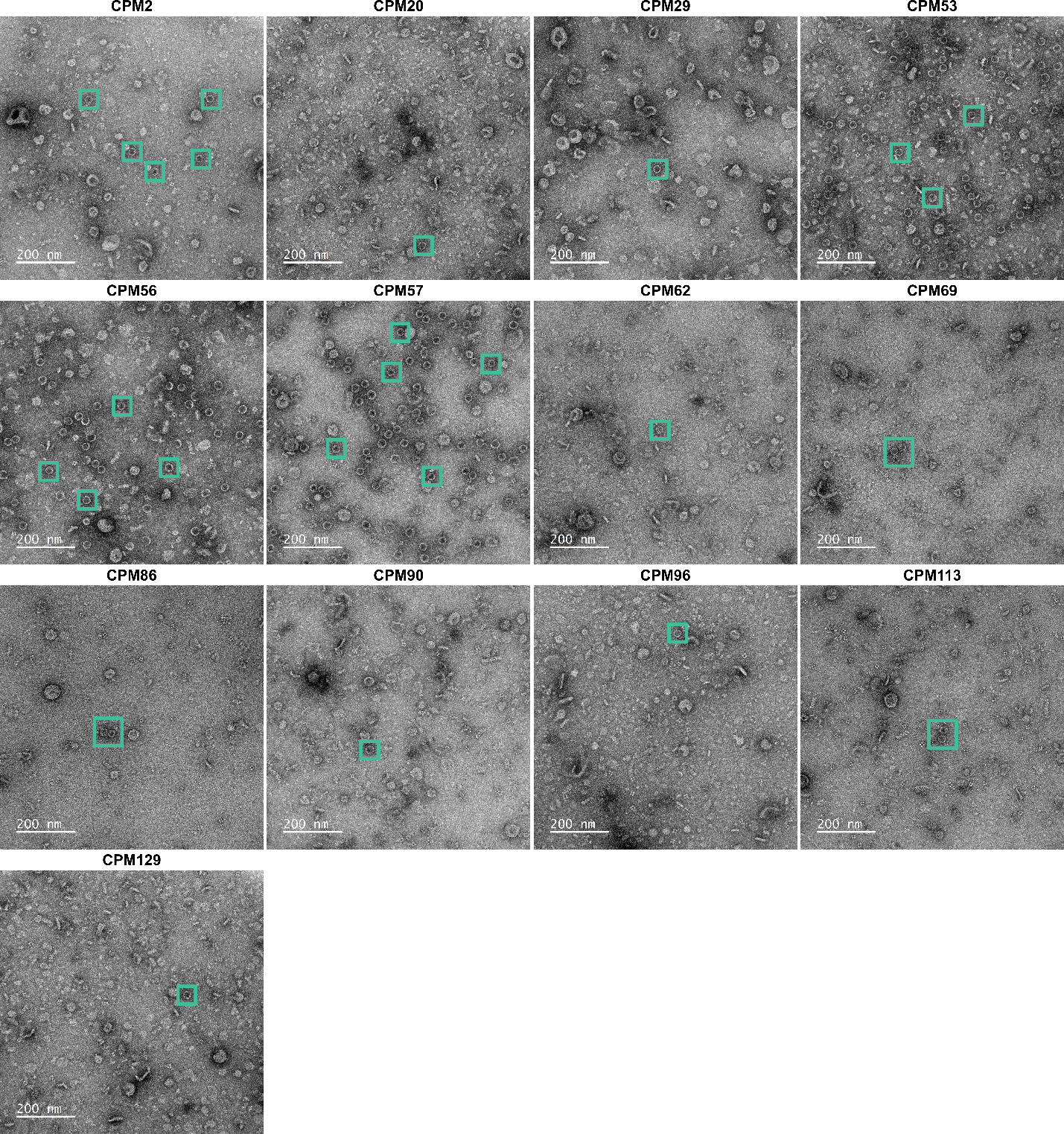

***Supplementary Fig. 4. TEM micrographs of selected permutants for individual assembly assays.*** VLP particles are boxed in mint. Samples were stained with uranyl acetate and imaged under the 50,000X magnification rate. CPM53, CPM56 & CPM57 were identified as assembly-competent permutants, CPM2 was classified as the intermediate permutant, and other permutants exhibited assembly deficiencies.

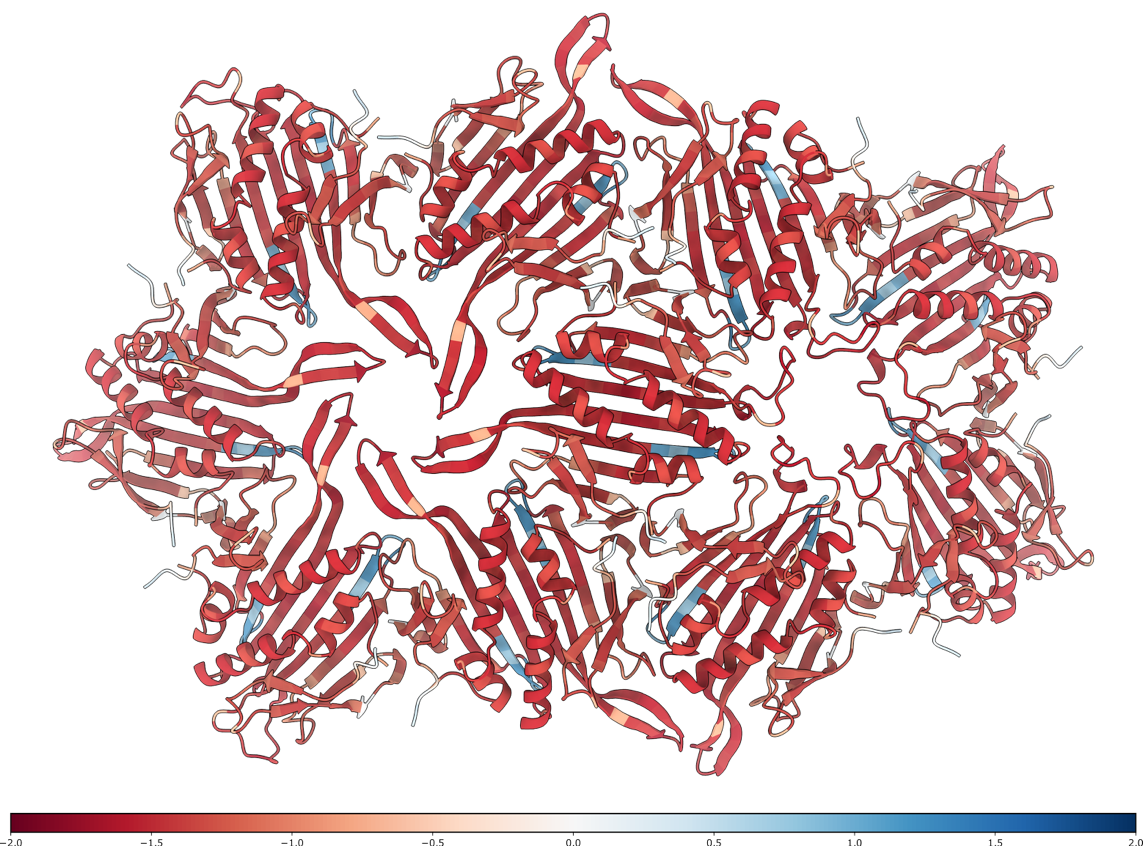

***Supplementary Fig. 5. MS2 VLP hexagonal (left) and pentagonal (right) facets colored by the circular permutation AFS.*** The exterior protein shell is at the front and the interior side is at the back (PDB ID: 1ZDI).

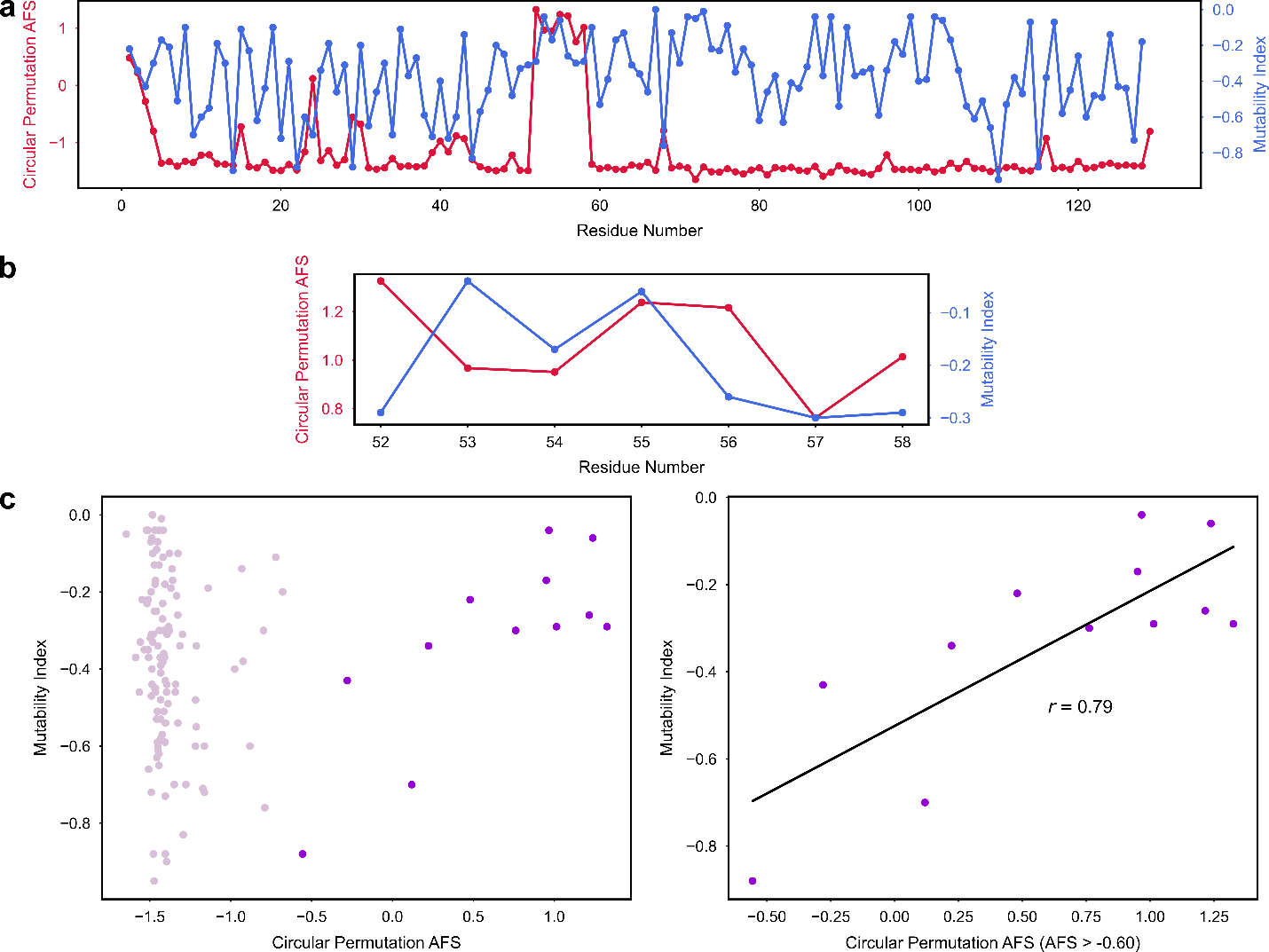

***Supplementary Fig. 6. Comparison between MS2 CP circular permutation AFS and mutability index.*** **a**, Circular permutation AFS and mutability index plotted by residue numbers. **b**, Comparison between circular permutation AFS and mutability index at the circular permutation super tolerant region. **c**, Circular permutation AFS and mutability index plotted as scatter plots. Residues with AFS > -0.6 (dark violet) were selected and their circular permutation AFS correlated with their mutability index with a Pearson correlation coefficient of 0.79.

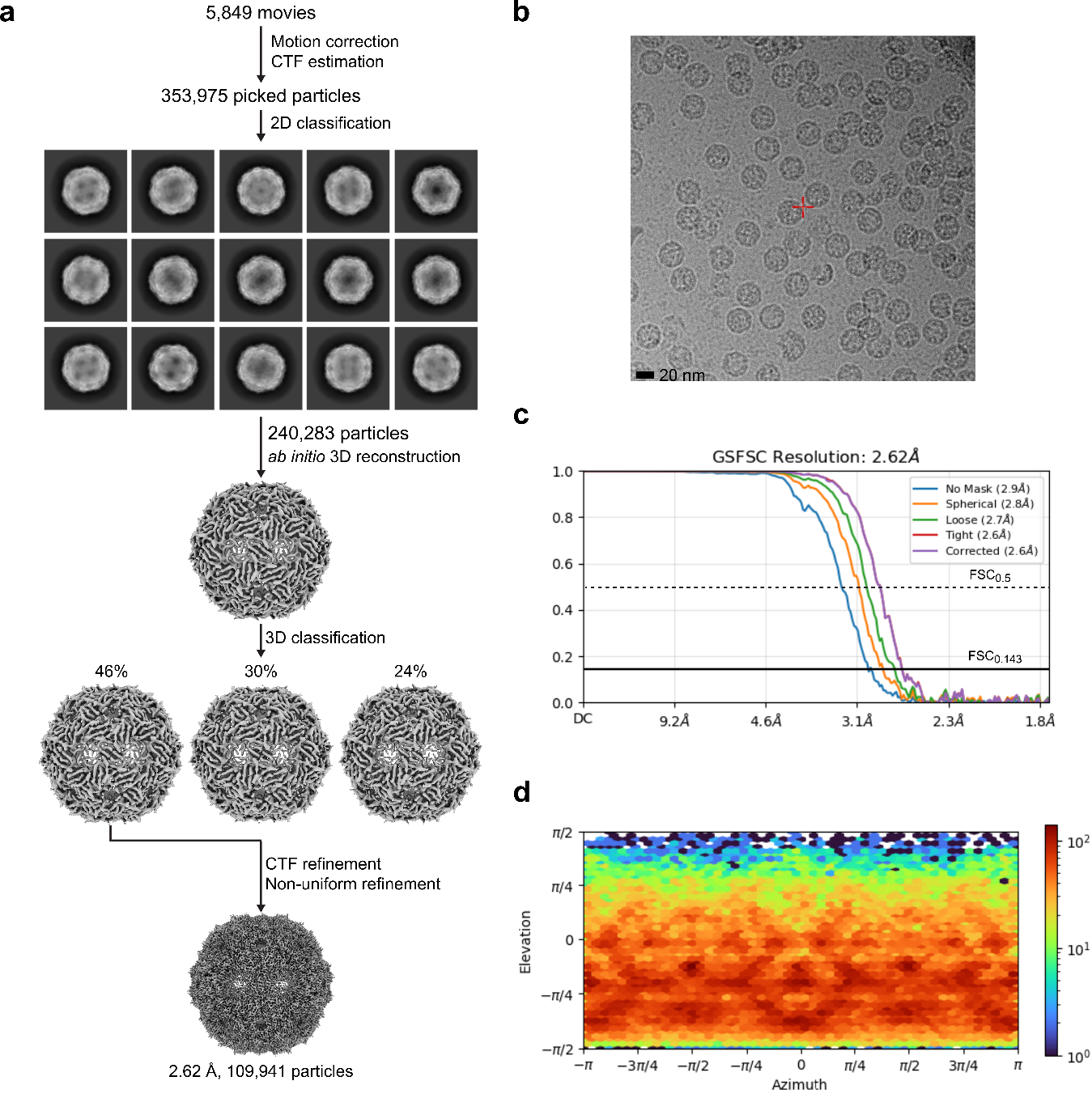

***Supplementary Fig. 7. Cryo-EM analysis of CPM58 VLP.* a**, Diagram of cryo-EM processing. **b**, A representative cryo-EM micrograph of CPM58 VLP. **c**, Gold Standard Fourier Shell Correlation (GSFSC) correlation plot of CPM58 VLP cryo-EM map. **d**, Viewing direction distribution plot of CPM58 VLP.
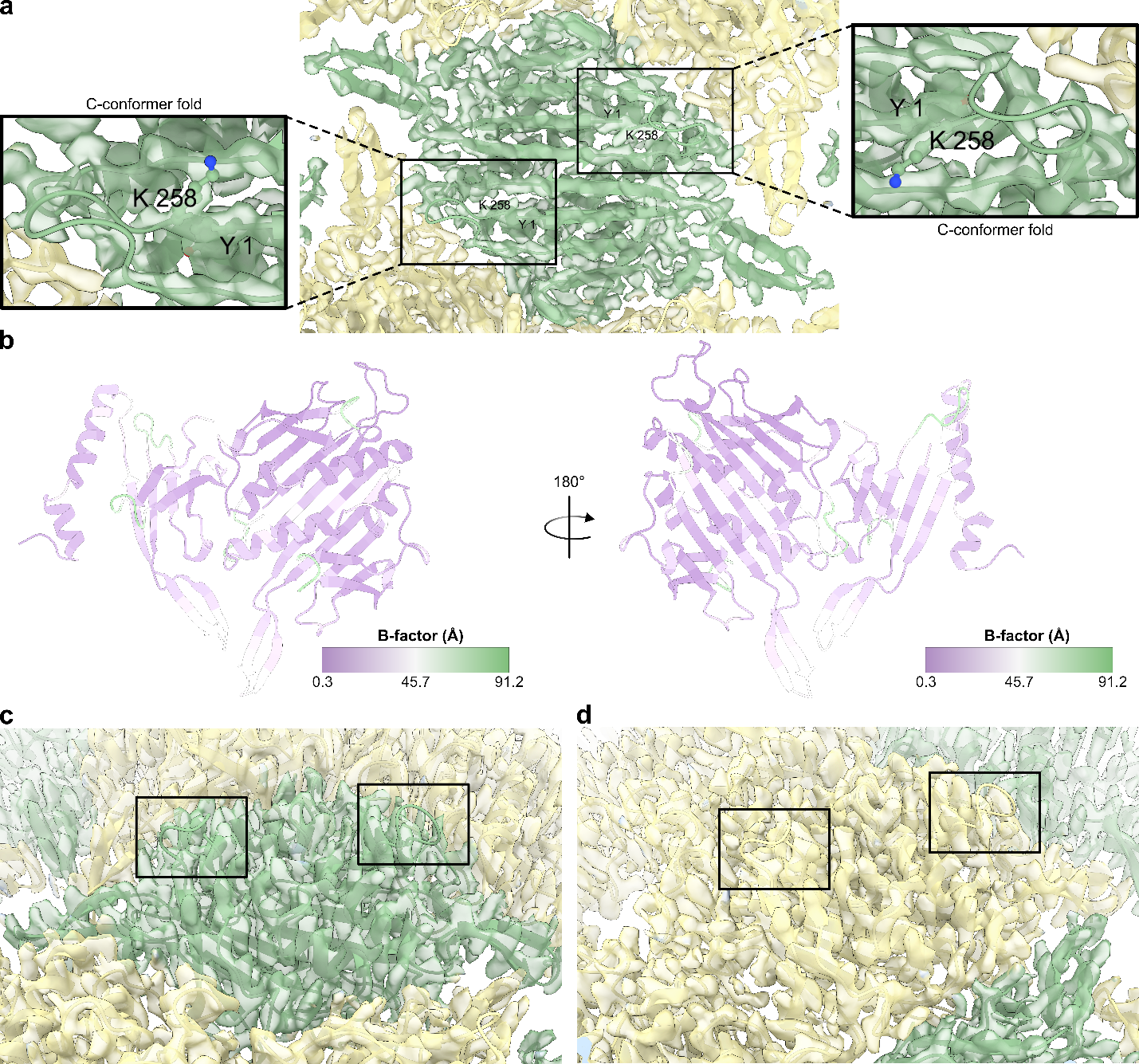

***Supplementary Fig. 8. Flexible structures in the CPM58 VLP subunits.*** **a**, Zoomed-in view of the cryo-EM density map around the symmetric subunit. The symmetric subunit was modeled as separate permuted CP chains without the intact CP sequence of the dimeric subunit. Residues corresponding to K57 and Y58 in the WT MS2 CP sequence are labeled and highlighted in ball-stick. **b**, CPM58 subunit models colored according to the B-factor pattern. **c**,**d**, Zoomed-in views of the cryo-EM density map around AB-loops (boxed) in the symmetric (**c**) and asymmetric (**d**) subunits.

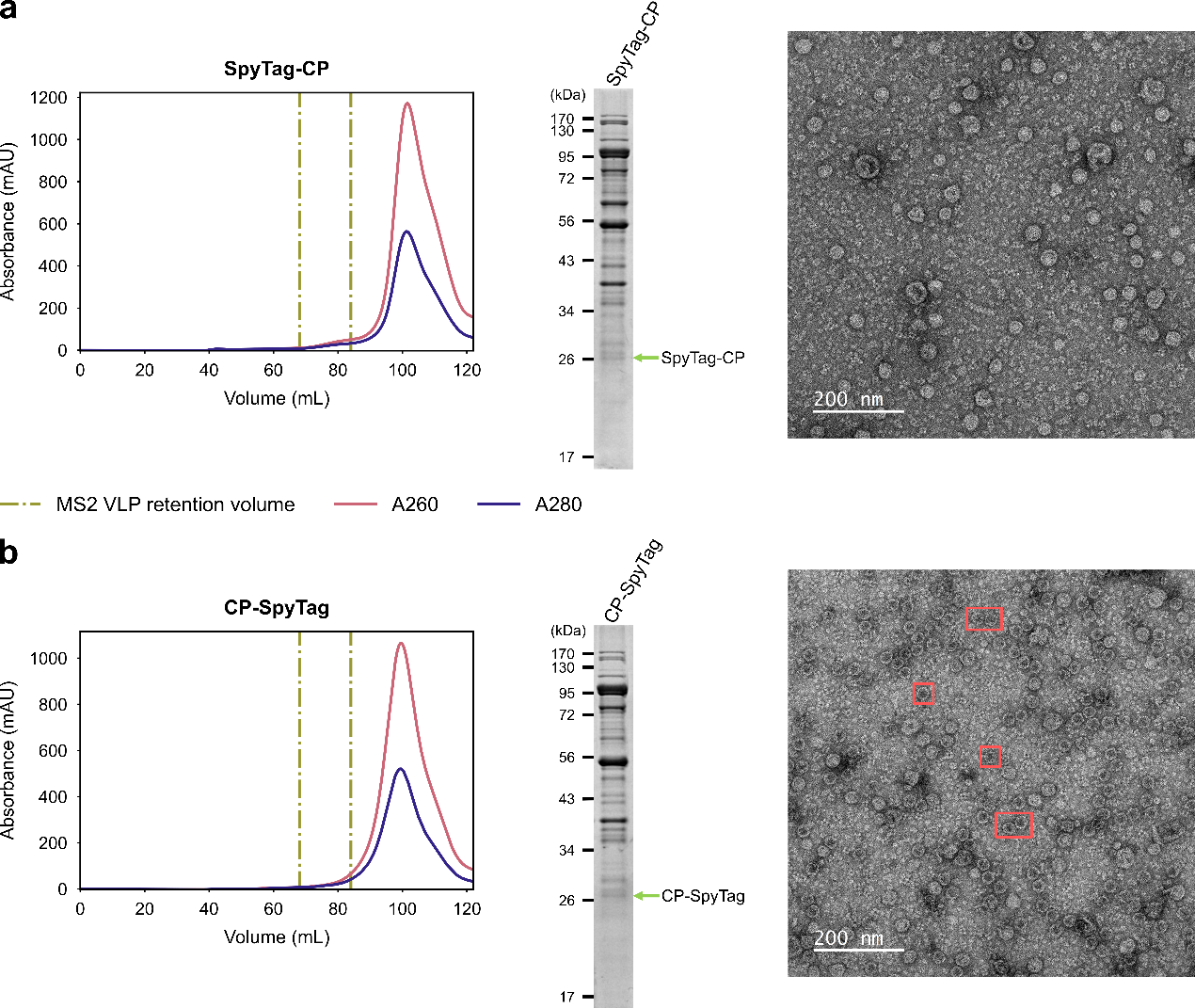

***Supplementary Fig. 9. Characterization of SpyTag-CP and CP-SpyTag VLPs.* a**,**b**, Purifications and characterizations of SpyTag-CP (**a**) and CP-SpyTag (**b**) VLPs by SEC, SDS-PAGE, and negative stain TEM. Classic VLP particles in TEM micrographs are boxed in red. No VLP particles were observed upon SpyTag-CP VLP purification. CP-SpyTag construct yielded in a low VLP titer.

***Supplementary Table 1. DLS measurements of MS2 VLPs***

| **variant** | **no peptide fusion** | | **permutant-SpyTag** | | **permutant-SpyTag/**  **SpyCatcher-sfGFP** | |
| --- | --- | --- | --- | --- | --- | --- |
|  | diameter (nm) | standard deviation (nm) | diameter (nm) | standard deviation (nm) | diameter (nm) | standard deviation (nm) |
| WT CP | 27.6 | 1.1 | -- | -- | -- | -- |
| CP-CP | 25.2 | 0.5 | -- | -- | -- | -- |
| CPM2 | 25.9 | 0.5 | 23.9 | 2.1 | -- | -- |
| CPM52 | 24.4 | 0.9 | 29.2 | 0.3 | 25.8 | 1.2 |
| CPM53 | 26.4 | 1.2 | 25.8 | 1.3 | 26.7 | 1.1 |
| CPM54 | 24.9 | 0.1 | 25.7 | 0.9 | 26.1 | 0.8 |
| CPM55 | 24.9 | 0.6 | 31.8 | 0.5 | -- | -- |
| CPM56 | 24.7 | 0.5 | 24.5 | 0.8 | 26.7 | 1.2 |
| CPM57 | 26.5 | 1.3 | 26.5 | 0.6 | 25.1 | 0.7 |
| CPM58 | 25.2 | 1.3 | 25.4 | 0.1 | 25.9 | 0.7 |

***Supplementary Table 2. Cryo-EM data collection, refinement and validation statistics.***

|  | **CPM58**  (EMDB-70484)  (PDB 9OH5) |
| --- | --- |
| **Data collection and processing** |  |
| Magnification | 130,000 |
| Voltage (kV) | 200 |
| Electron exposure (e^–^/Å^2^) | 60 |
| Defocus range (μm) | -0.6 to -1.6 |
| Pixel size (Å) | 0.894 |
| Symmetry imposed | I |
| Initial particle images (no.) | 353,975 |
| Final particle images (no.) | 109,941 |
| Map resolution (Å)  FSC threshold | 2.6  0.143 |
| Map sharpening *B* factor (Å^2^) | -113.6 |
| **Refinement** |  |
| Initial model used (PDB code) | 1ZDI |
| Model composition  Non-hydrogen atoms  Protein residues  Nucleic acid residues  Ligands  Refinement (Phenix)  Map correlation coefficient  (whole unit cell)  Map correlation coefficient (around atoms)  R.m.s. deviations  Bond lengths (Å)  Bond angles (°) | 2,894  387  0  0  0.43  0.80  0.002  0.552 |
| Validation  MolProbity score  Clashscore  Poor rotamers (%) | 1.35  2.94  1.57 |
| Ramachandran plot  Favored (%)  Allowed (%)  Disallowed (%) | 97.38  2.62  0.0 |
